# Activating mutations in FGFR3 are associated with clonal expansion events and high *de novo* rates in the male germline

**DOI:** 10.1101/2022.07.31.502216

**Authors:** Sofia Moura, Ingrid Hartl, Atena Yasari, Veronika Brumovska, Yasmin Striedner, Marina Bishara, Theresa Mair, Thomas Ebner, Gerhard J. Schütz, Eva Sevcsik, Irene Tiemann-Boege

## Abstract

Delayed fatherhood results in a higher risk to inherit a new germline mutation that might result in a congenital disorder in the offspring. In particular, some *FGFR3* mutations increase in frequency with age, but there are still a large number of uncharacterized *FGFR3* mutations that could be expanding in the male germline with potentially early or late-onset effects in the offspring. Here, we investigated the mutation frequency in the DNA of human testis and sperm and the activation state of the expressed mutant protein of eight different *FGFR3* variants categorized by ClinVar as deleterious, benign, or not reported. Overall, the ligand-independent activation of the mutant protein resulted in a increased number of mutant sperm; although, strong activating mutations did not necessarily result in the highest frequencies. Moreover, only two mutants c.952G>A and c.1620C>A showed an increase with the donor’s age; the latter also forming larger clonal expansions in the testis. We also showed that the prediction of deleteriousness of a mutation is not always accurate, and similar *in silico* scores can reflect either a gain-of-function or loss-of-function. Our approach led to the discovery of two novel variants c.1261G>A and c.952G>A to have promiscuous *FGFR3* activation and increased mutation frequencies in the male germline. The large fraction of donors with mutations suggests a high *de novo* rate potentially explained by a selective advantage before the maturation of the male germline. This sequence-function study provides important data for the evaluation and interpretation of variants with relevant clinical implications.

## Introduction

While the long-term consequences of advanced maternal age on offspring are well-established (Tarín et al., 1998), the higher risk of progeny of older fathers inheriting a congenital disorder is not so well-known. The ongoing germline cell division occurring throughout the male lifetime leads to the accumulation of *de novo* mutations (DNMs) in the male germline (Campbell et al., 2015; Kong et al., 2012). DNMs occur rather rarely and are lost by genetic drift. However, a special class of mutations has been measured at low frequencies (10^−6^ to 10^−4^) in the male germline, orders of magnitude higher than the estimated average human genome mutation rate per cell division per generation at any given genomic position (∼1.1×10^−8^) (Francioli et al., 2015; Kong et al., 2012). These mutations affect cell growth and proliferation, act by positive selection and as a result, increase in number with the age of the male germline and might have a higher chance to be passed on to the progeny by older fathers (reviewed in (Arnheim and Calabrese, 2016, 2009; Goriely et al., 2009; Goriely and Wilkie, 2012)).

Such driver or selfish mutations are described in tumors (Jamal-Hanjani et al., 2017; Turajlic et al., 2018; Yates et al., 2015). In the male germline they have been observed in genes involved in the RAS-MAKP signaling pathway (e.g., *FGFR2, FGFR3, RET, HRAS, PTPN11, KRAS BRAF, CBL, MAP2K1, MAP2K2, RAF1, and SOS1)* (Choi et al., 2012; Goriely et al., 2009; Green et al., 2010; Maher et al., 2018, 2016; Qin et al., 2007; Shinde et al., 2013). Some missense, gain-of-function mutations in these genes are enriched in sperm and testis and result in autosomal dominant genetic disorders. They occur exclusively in the male germline and older men have a higher probability of having an affected child than younger males - known as the paternal age-effect (PAE) described already decades ago (Crow, 2012, 2000; Risch et al., 1987), or as RAMP mutations (Recurrent, Autosomal dominant, Male biased, and Paternal age effect) (Arnheim and Calabrese, 2016) or selfish mutations (Goriely and Wilkie, 2012).

To date, the best characterized congenital disorders associated with a PAE are Apert (Glaser et al., 2003; Risch et al., 1987; Tolarova et al., 1997), Crouzon (Risch et al., 1987), Pfeiffer (Risch et al., 1987), Costello (Zampino et al., 2007) and Noonan syndromes (Tartaglia et al., 2004; Yoon et al., 2013), Multiple endocrine neoplasia type 2B (Choi et al., 2012), Muenke craniosynostosis (Rannan-Eliya et al., 2004), Hypochondroplasia (Goriely and Wilkie, 2012), Thanatophoric Dysplasias (Goriely et al., 2009), and Achondroplasia (Orioli et al., 1995; Shinde et al., 2013; Tiemann-Boege et al., 2002), all caused by the aforementioned genes. Not only PAE-mutations causing early-onset diseases (e.g., rasopathies or skeletal dysplasias) have been described to increase with paternal age. Reports have also documented an increased frequency with paternal age of late-onset neurological and behavioral disorders including autism (Frans et al., 2013; Hultman et al., 2011; Kong et al., 2012), schizophrenia (Svensson et al., 2012), and bipolar disorders (Frans et al., 2008; Grigoroiu-Serbanescu et al., 2012). Also, breast cancer or cancers in the nervous system are hypothesized to correlate with paternal age (Hemminki et al., 1999; Hemminki and Kyyrönen, 1999), and reviewed in (Acuna-Hidalgo et al., 2016; Paul and Robaire, 2013; Sharma et al., 2015).

Several studies have demonstrated that the specific molecular mechanisms of certain PAE disorders are explained by changes in the function of the mutant protein expressed in the male germline. In particular, driver mutations in genes of receptor tyrosine kinases (RTKs), such as *FGFR3*, modify the signaling of the receptor by promoting ligand-independent activation (Bellus et al., 2000; He et al., 2012; Krejci et al., 2008; Sarabipour and Hristova, 2016). This promiscuous receptor activation occurring in the self-renewing spermatogonial stem cells (SrAp) promotes a positive selective advantage compared to the neighboring cells. As a result of continuous cell divisions, a high mutation burden will accrue in the testis over time forming clusters or sub-clonal populations and a higher number of mutant sperm in older donors, as described in *FGFR3* for p.G380R (Shinde et al., 2013; Tiemann-Boege et al., 2002); and different mutations in codon 650 (Goriely et al., 2009) and p.R669G (Maher et al., 2018). Driver mutations form clonal subpopulations varying in size, and all found in different anatomical locations of the testes (Arnheim and Calabrese, 2016; Maher et al., 2018; Shinde et al., 2013).

Despite the importance of driver mutations in the male germline with their high incidence, the increased frequency with paternal age, and early- or late-onset effects, we know very little about this mutagenic mechanism. Further, with the explosion of sequencing data of somatic tissues in clinical settings (e.g., COSMIC), a large number of mutations are being reported for *FGFR3*. Many have unknown effects but could be expanding with paternal age with phenotypes potentially leading to early- or late-onset disorders in children of older men. However, the impact of genetic changes on the activation state and the resulting sub-clonal expansion in the male germline of different *FGFR3* variants is not well-understood. There is only one analysis of this kind for codon 650, an important tyrosine kinase (TK) activation site (Goriely et al., 2009). In this work, it was shown that strongly activating mutations often associated with tumors (p.K650M), were found at lower frequencies in sperm than milder activating mutations (Goriely et al., 2009).

As such, identifying prospective driver mutations increasing in number with the aging male germline has been an intensive target of research, but a tricky task given the ultra-low frequency of these events (Maher et al., 2018; Salazar et al., 2022). Advances in next-generation sequencing (NGS) technologies have enabled the identification of mutations with high resolution. In particular, identifying prospective driver mutation in the testes (Maher et al., 2018) or sperm (Salazar et al., 2022) using ultra-deep sequencing methods have identified a series of mutations in the RTK-RAS pathway expanding in the male germline. Especially, duplex sequencing has the sensitivity to capture mutations occurring as low as 10^−8^ to 10^−6^ (Abascal et al., 2021; Hoang et al., 2016; Kennedy et al., 2014; Kinde et al., 2011; Schmitt et al., 2012) and has identified rare variants in *FGFR3* of sperm DNA occurring at higher than expected frequencies (Salazar et al., 2022). However, despite the promising sensitivity of duplex sequencing, in practice, it still lacks the throughput to screen enough genomes needed to assess larger sample sizes.

To gain further insights into the accumulation of driver mutations with age, their effect on spermatogenesis, and their signaling properties, we analyzed eight *FGFR3* missense substitutions in sperm DNA of donors ranging in age from 23 to 59 years of age using droplet digital PCR (ddPCR). Five of the variants have been associated with congenital disorders; three of them were predicted *in silico* to have a strong deleterious effect on the protein structure (CADD) and described as deleterious (ClinVar), and two had uncertain or likely benign effects. Three variants had never been characterized nor have been reported, but were recently found at higher frequencies in sperm DNA (Salazar et al., 2022). We also studied the spatial distribution of three variants in human testis of a 73-year-old donor. We identified two paternal age effect variants: c.952G>A and c.1620C>A, with the latter promoting clonal expansion in the testis. Activation studies in HeLa cells revealed an increased signaling activation (even in the absence of ligand) for six mutants. The remaining sites either showed no difference from the wild-type or decreased signaling. Our data further stresses the importance of examining *FGFR3* variants expanding in the male germline with important clinical implications to better understand the biological and biochemical effects of potentially causal variants of disease.

## Results

### Measuring low-frequency variants with ddPCR

First, we determined the accuracy and reproducibility of ddPCR as a detection method for each of the eight specific mutations. For this purpose, we performed serial dilution experiments with positive controls (cell line encoding c.1620C>A or sequenced-confirmed variant plasmids for the remaining target loci) spiked into wild-type (WT) human genomic DNA at different ratios (Figure 1A). We observed a good correspondence between the input ratio of mutant to wild-type copies with respect to the observed ratio, with a higher variation for the lower copy input which can be explained by e.g., larger deviations by the Poisson distribution of rare events. The strong correlation between replicates for each dilution step (Pearson’s R, ρ, ranging from 0.794 to 0.997) indicated the accuracy and reproducibility of ddPCR for the eight mutations between a dynamic range of 10^−1^ to 10^−4^.

**Figure 1:**
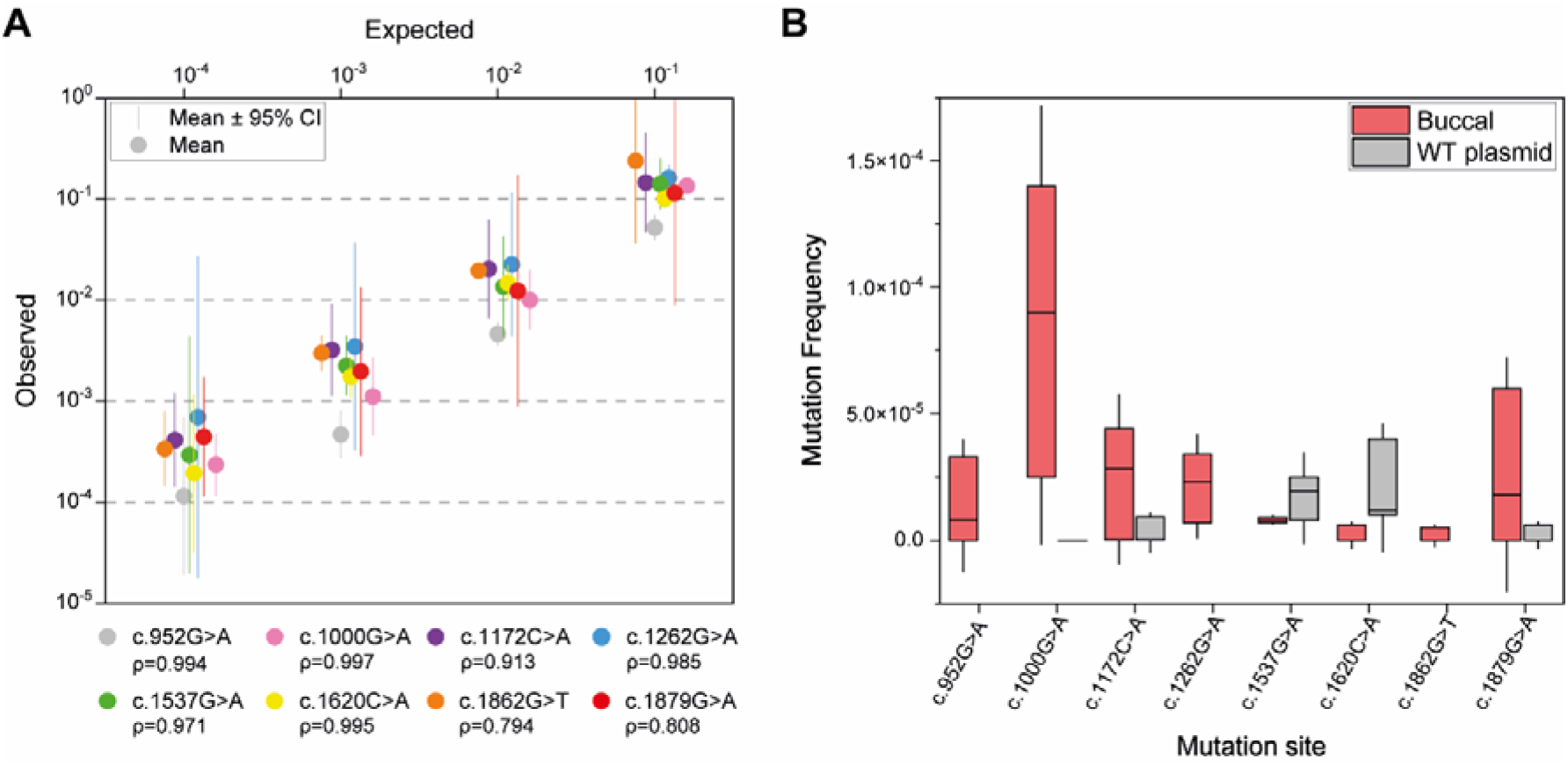
Control testing. **(A)** Spike-in serial dilutions for the eight target sites and their respective Pearson’s correlation coefficient (ρ). We measured three biological replicates per dilution step with a mixture of WT (human genomic DNA) and variant templates (Coriell Cell line or variant plasmids) according to each dilution step (1:10 to 1:10,000, 10-fold steps). Dilutions contained either ∼20,000, ∼36,000, or ∼72,000 genomes of wild-type (human genomic DNA) and variant templates distributed in 1, 2, or 4 reactions, respectively such that there was on average one DNA template per droplet. **(B)** Variant detection in two controls hypothesized to be mutation-free: buccal epithelial cell DNA and WT plasmid. We measured three biological replicates (n = ∼300,000 copies per replicate with four or three reactions with ∼72,000 or ∼100,000 copies per reaction, respectively) for each variant site for buccal and/or plasmid samples, except for site c.1537G>A with 4 biological replicates for the plasmid (Supplementary Table S1).

Second, we also assessed the false-positive rate (mutant counts due to technical artifacts) using negative controls. The negative controls we used were sequence confirmed plasmids and/or DNA from non-invasive somatic tissue (e.g., buccal epithelial cells reported to have no *FGFR3* expression (Uhlen et al., 2019; Uhlén et al., 2015)). Specifically, we screened the same copy number (∼300,000 copies) of buccal cells or plasmids as for sperm and testis DNA (Figure 1B; Table S1). We measured a lower median VAF (non-significant) for plasmids and/or buccal cells than for sperm samples (except for c.1537G>A, see Supplementary Tables S1 and S2); although, the VAF for the negative controls varied extensively between sites or even replicates. In general, the WT plasmid measured a median VAF= 0 for three out of five variants (c.1000G>A, c.1879G>A, and c.1172C>A); whereas the buccal DNA median VAF was 0 for 2 out of 8 sites (c.1620C>A and c.1862G>T) and ranged between 9.0×10^−5^ to 8.0×10^−6^ for the other sites (Supplementary Table S1). The VAF value of the negative controls was independent of the type of base substitution, but in general, transversions had a lower VAF for control samples than sperm (Supplementary Table S2), in spite of all eight substitutions being also common DNA lesions with C>A/G>T attributed to guanine oxidation and G>A/C>T transitions to cytosine deamination (Arbeithuber et al., 2016).

The selection of reliable negative controls without mutations remains highly challenging. *E. coli* has a high-fidelity replication machinery with an error rate as low as 5.4×10^−10^ per base pair per replication (Drake et al., 1998). In theory, this low error replication rate would make sequence-confirmed wild-type plasmids a hypothetically mutation-free control. In our measurements, this was not always the case with oxidation and heat exposure possibly contributing to the number of mutations to levels of 10^−6^ to 10^−5^ from these common DNA lesions (Arbeithuber et al., 2016; Jee et al., 2016). Also note that the VAF measured for the buccal epithelial cells (∼5×10^−6^) was also not a mutation-free control and measured frequencies within the range reported for somatic tissue, e.g., blood (∼10^−6^) (Abascal et al., 2021). This emphasizes that somatic tissue or WT plasmids might not be mutation-free and thus, are not reliable negative controls. Alternatively, these results might also represent background noise of the ddPCR method.

In order to investigate this further, we also used the number of sperm samples rendering no mutant counts as a proxy for the sensitivity of each assay. As such, for all eight variants, we counted the number of WT genomes in samples selected without any mutant counts, which translated to a sensitivity well below <10^−7^ (e.g., for c.1862G>T no mutants were counted for 66% of the measured sperm samples which can be translated to 0 mutations in 10^7^ WT genomes or a VAF below ∼7×10^−8^ as shown in Table S3). For a more detailed discussion on the sensitivity of the assay please refer to Supplementary Methods.

### Mutations in sperm DNA

Here, we further investigated in a total of 177 sperm donors (aged 23-59) the mutation frequency of the eight *FGFR3* variants (Table S4). We selected our candidate mutations (Table 1) based on their association with phenotypes, clinical significance (pathogenic, likely benign, uncertain), predicted deleteriousness (CADD score, (Kircher et al., 2014)), gnomAD and/or COSMIC reports, and lastly, according to recent elevated VAF reports in younger and older donors (Salazar et al., 2022). Three out of eight variants were classified in ClinVar as deleterious/pathogenic, two were benign or uncertain, yet had a very high CADD score and the remaining variants had an unknown ClinVar categorization, but had a high CADD score and were reported in gnomAD and/or COSMIC. Seven of our candidate mutations have been reported to occur in the DNA of young and/or old donors at frequencies ∼10^−5^ to 10^−4^ with duplex sequencing (Salazar et al., 2022).

**Table 1:**
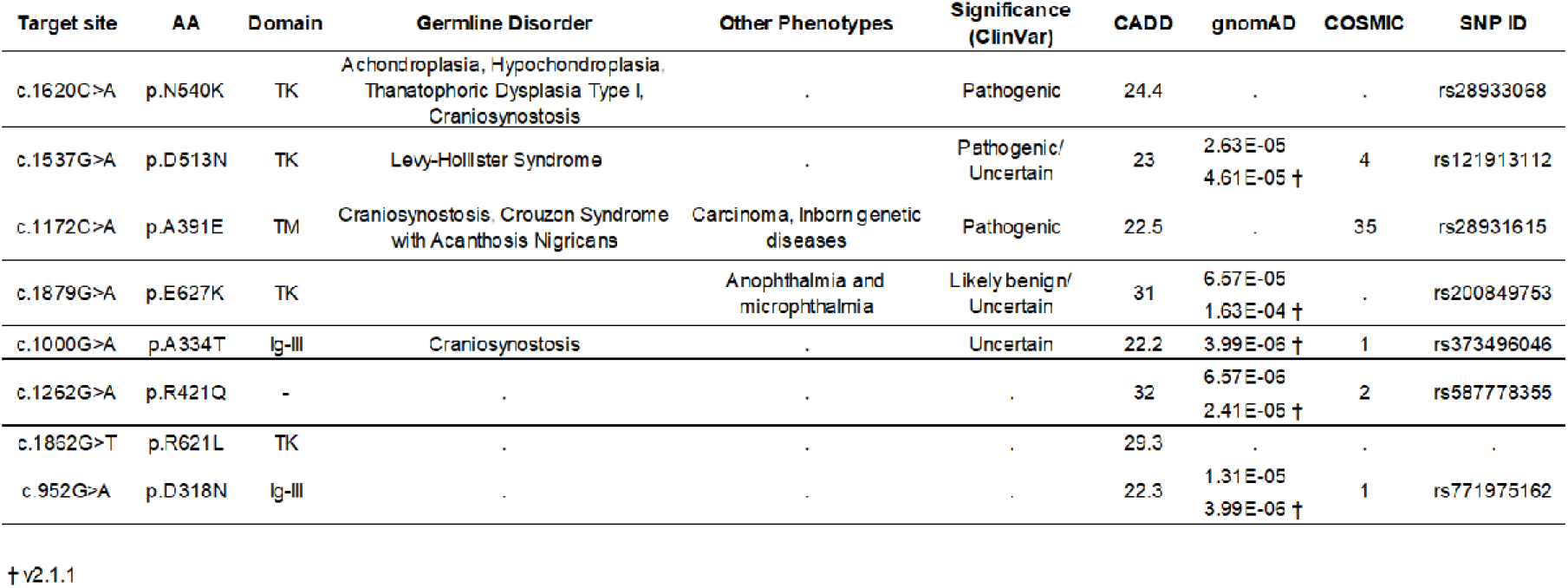
Target *FGFR3* variants and associated information. Variants are organized by significance (ClinVar database) followed by the CADD score (Kircher et al., 2014). Phenotype data were retrieved from ClinVar. CADD (Combined Annotation Dependent Depletion) score is defined according to GRCh38-v1.6 and annotates the level of deleteriousness. All gnomAD information was retrieved from v3.1.2 except those indicated (†) (Karczewski et al., 2020). COSMIC (Catalogue Of Somatic Mutations In Cancer) data was based on version 94 (Tate et al., 2019). All data was called based on transcript ENST00000440486, *FGFR3* isoform IIIc, and was last updated on the 16^th^ of July, 2022.

We first assessed if there was an increase in mutations with the donor’s age. Our Spearman’s correlation test for each mutation site (Figure 2A), showed a significant positive correlation in variant frequency with age only for two sites: c.952G>A (R=0.21, p=0.0497) and c.1620C>A (R=0.34, p=0.00029). The third site (c.1000G>A; p.A334T) also showed a positive correlation of VAF with age, but bordered on significance (R=0.2, p=0.06). Note that the VAF was independent of other factors such as sperm diagnosis or sperm count (Million sperms/mL) and no correlation was observed when testing the 177 screened donors (Table S5). We detected the strongest correlation between age and variant frequency (average VAF: 1.1×10^−5^ in 108 donors, Table 2) for variant c.1620C>A, with the highest VAF of 1×10^−4^ and the lowest at 0 (Table S4). Further age sub-categorization of the sperm data revealed a significant difference between the VAF of younger (≤30 years) and middle (31-44 years) groups (p=0.0002), and also between the younger and older (≥45 years) age categories (p=0.0031; Figure 2B). For the majority of the donors (63%) we counted mutations for this variant, but almost one-third of the donors had a VAF of 0 (Figure 2C). Overall, the measured frequencies in sperm DNA for the c.1620C>A variant were in accordance with the VAF recently published (Salazar et al., 2022) and reported in Table 2.

**Table 2:** Average and median variant allele frequencies estimated for the different *FGFR3* variants in sperm DNA. AA: Amino Acid. VAF: Variant Allele Frequency. IQR: Interquartile range.*Data from duplex sequencing (Salazar et al., 2022).

| Loci | AA | Sperm |  |  |  |  | VAF* |
| --- | --- | --- | --- | --- | --- | --- | --- |
|  |  | n | Mean VAF | Standard Deviation | Median VAF | IQR |  |
| c.1620C>A | p.N540K | 108 | $1.1 \times 10^{-5}$ | $\pm 1.6 \times 10^{-5}$ | $5.0 \times 10^{-6}$ | $\pm 1.8 \times 10^{-5}$ | $1.9 \times 10^{-5}$ |
| c.1537G>A | p.D513N | 91 | $9.4 \times 10^{-6}$ | $\pm 1.7 \times 10^{-5}$ | $5.0 \times 10^{-6}$ | $\pm 1.0 \times 10^{-5}$ | $2.8 \times 10^{-5}$ |
| c.1172C>A | p.A391E | 79 | $8.6 \times 10^{-6}$ | $\pm 1.8 \times 10^{-5}$ | 0 | $\pm 8.0 \times 10^{-6}$ | . |
| c.1879G>A | p.E627K | 79 | $1.5 \times 10^{-5}$ | $\pm 1.3 \times 10^{-5}$ | $1.1 \times 10^{-5}$ | $\pm 1.4 \times 10^{-5}$ | $5.3 \times 10^{-5}$ |
| c.1000G>A | p.A334T | 90 | $9.1 \times 10^{-5}$ | $\pm 7.4 \times 10^{-5}$ | $8.5 \times 10^{-5}$ | $\pm 1.2 \times 10^{-4}$ | $2.0 \times 10^{-4}$ |
| c.1262G>A | p.R421Q | 99 | $1.7 \times 10^{-5}$ | $\pm 2.1 \times 10^{-5}$ | $1.1 \times 10^{-5}$ | $\pm 2.0 \times 10^{-5}$ | $1.9 \times 10^{-5}$ |
| c.1862G>T | p.R621L | 93 | $2.7 \times 10^{-6}$ | $\pm 4.5 \times 10^{-6}$ | 0 | $\pm 5.0 \times 10^{-6}$ | $1.9 \times 10^{-5}$ |
| c.952G>A | p.D318N | 91 | $2.2 \times 10^{-5}$ | $\pm 2.4 \times 10^{-5}$ | $1.5 \times 10^{-5}$ | $\pm 2.0 \times 10^{-5}$ | $1.9 \times 10^{-5}$ |

**Figure 2:**
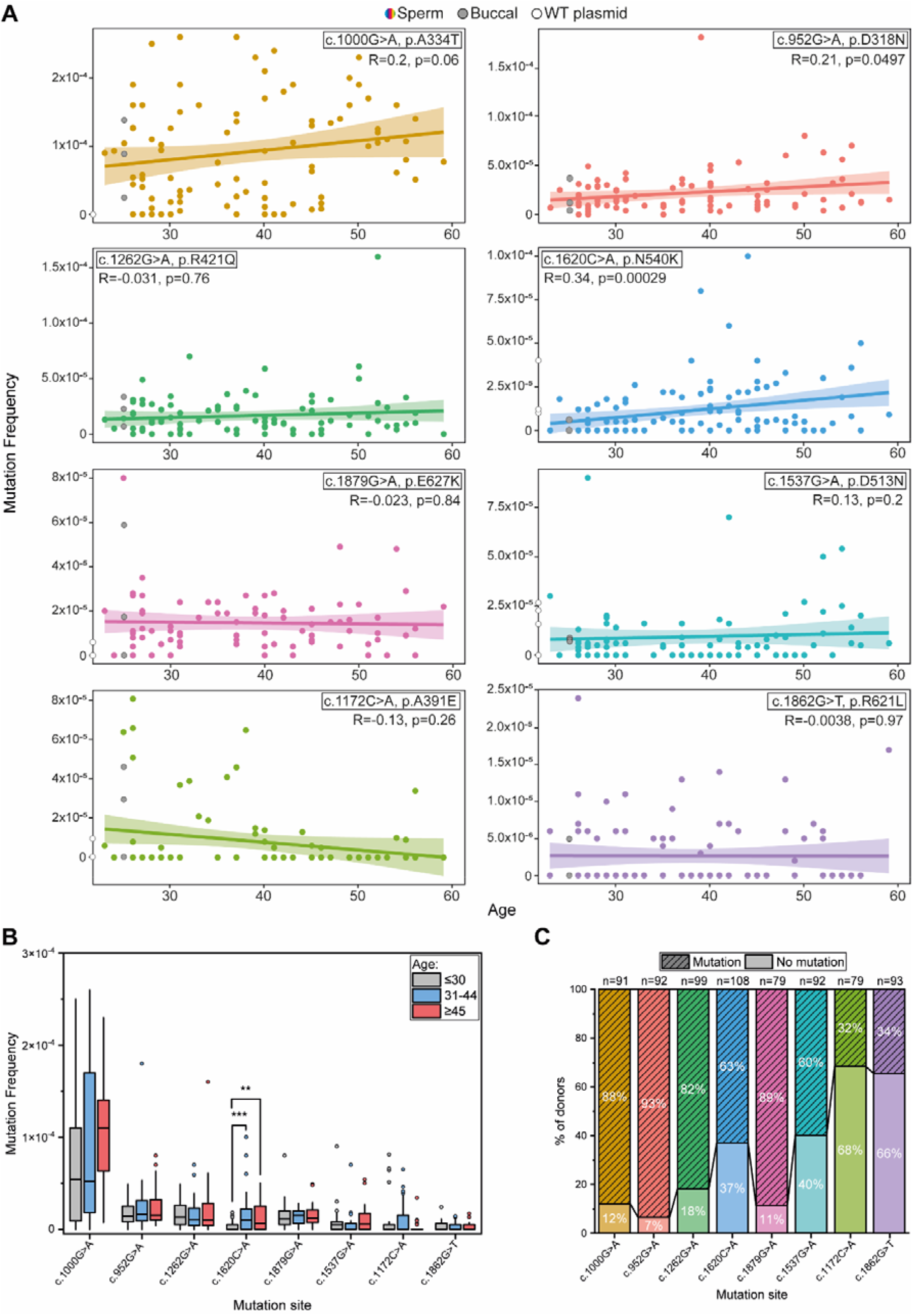
Analysis of eight *FGFR3* mutations in sperm DNA. **(A)** Mutation frequency of *FGFR3* variants measured in 177 sperm DNA donors. Spearman’s correlation test was performed for each individual mutation. For comparison, both negative controls, buccal cell DNA (25 years old; grey) and WT plasmid (white circles on the y-axis), were also plotted. **(B)** Mutation distribution in three age groups: younger (≤30), middle (31-44), and older (≥45) in sperm DNA for our candidate mutations. Tests for significant differences between the three age categories were performed with the Kruskal Wallis test; P-values were estimated using the Mann-Whitney-U test annotated as p≤0.01 (**) and p≤0.001 (***). Mutation sites are organized by highest to lowest mutation frequency IQR values. **(C)** Percentage of donors that do and do not harbor mutations for the eight *FGFR3* loci. The number of total donors screened for each locus is represented (n) at the top of each bar column. Percentages are rounded with no decimal number. **(A and C)** Coding sequence substitutions are color coded.

In comparison, the two other variants categorized as pathogenic by ClinVar (c.1537G>A; p.D513N and c.1172C>A; p.A391E) did not show an increased VAF with age (Figure 2A). However, we observed for variant c.1537G>A that also 60% of the 92 screened donors harbored this mutation (mean VAF: 9.4 ×10^−6^; Figure 2C, Table 2). The VAF was similar across the different age categories and the Spearman’s test did not show a significant correlation between VAF and age (Figures 2A and 2B). It is possible that for this site the method is not sensitive enough and lacks the power to detect differences at a VAF <10^−5^ and thus a positive correlation is not measurable. However, given that no mutation was measured for 40% of the sperm samples, which can be translated to a sensitivity of <10^−7^ (see Supplementary Table S3), it is more likely that a different expansion mechanism could be taking place that does not fit the classical strong clonal expansion patterns with age as observed with variant c.1620C>A

Similarly, Spearman’s correlation test revealed no significant effect in VAF with age for variant c.1172C>A. The average VAF was 8.6×10^−6^ (Table 2) and mostly younger donors carried this mutation. Although this variant has the lowest deleterious score of the three pathogenic variants, it has the highest COSMIC hits in tumor samples of all eight variants. Notably, we only detected this variant in 32% of our donors (Figure 2C). Several reasons could explain this trend: 1) mutations are truly absent, 2) this variant occurs at a much lower frequency detectable only when screening more genomes, or 3) the few measured mutant counts are artifacts. Given that the measured VAF was as low as 10^−6^ and that the sum of all screened genomes without a mutation equal to a VAF of <10^−7^, the first two explanations are more likely. Thus, this variant is likely expanding at very low levels.

Next, the variants (c.1879G>A and c.1000G>A) categorized as likely benign and/or uncertain significance had different deleteriousness scores (31 and 22, respectively), but were found in a similar percentage in the different donors. Variant c.1879G>A was detected in 89% of our donors (n=79) at an average frequency of 1.5×10^−5^ but showed no correlation between VAF and age (Figure 2A) nor differences between the three age categories (Figure 2B). Variant c.1000G>A (p.A334T) had the highest VAF (9.1×10^−5^) of all eight variants with 88% of our sperm donors harboring mutations (Table 2, Figure 2C). Further, we observed a positive correlation between age and VAF, yet it was not significant (p= 0.06). We cannot explain this high number of mutations found in a large number of donors including the younger group (88%), but we hypothesize that this mutation is tolerated at higher frequencies, or is a common somatic mutation that occurred during development, or in the immature male germline affecting a higher proportion of cells.

### Characterization of novel variants

In this work, we also analyzed uncharacterized variants c.1262G>A (p.R421Q), c.1862G>T (p.R621L), and c.952G>A (p.D318N) that have a high predicted deleteriousness score, and have recently been reported at higher frequencies in sperm DNA (Salazar et al., 2022). Our screening revealed that 82% of our donors were positive for the c.1262G>A variant at an average VAF of 1.7×10^−5^ (Figure 2C and Table 2), similar to the levels reported in (Salazar et al., 2022). Our data showed neither a correlation between VAF and age (Figure 2A) nor a difference in frequency between the three age groups (Figure 2B).

Another interesting novel variant is c.1862G>T which has been reported in the literature with a different amino acid substitution at the same codon (c.1862G>A, p.R621H) as a pathogenic variant with a slightly higher CADD score (29.8) associated with CATSHL (Camptodactyly, Tall Stature, and Hearing Loss) Syndrome (Toydemir et al., 2006). Sperm DNA analysis revealed that similar to the c.1172C>A variant, there is no age correlation. Only 34% of our donors were carriers of this variant and it had the lowest average VAF of 2.7×10^−6^ (Figure 2 and Table 2). Lastly, the c.952G>A variant has one of the lowest predicted CADD scores but has been reported in both gnomAD and COSMIC databases. Interestingly, this variant showed a significant positive correlation in variant frequency with age with 93% of donors harboring mutations (mean frequency: 2.2×10^−5^; Figure 2 and Table 2).

### Spatial distribution of variants in aged human testis

To further test the clonal expansion in the male germline, we screened the spatial distribution of three variants in the human testis of a 73-year-old post-mortem donor (Figure 3F). Our selection of the screened variants was based on 1) the variants showing a positive correlation between age and VAF with a possible paternal age-effect (PAE) (c.1620C>A and potentially c.1000G>A) and 2) no clear increase with age and no association with any phenotypes, but a high VAF and a high CADD score (c.1262G>A).

**Figure 3:**
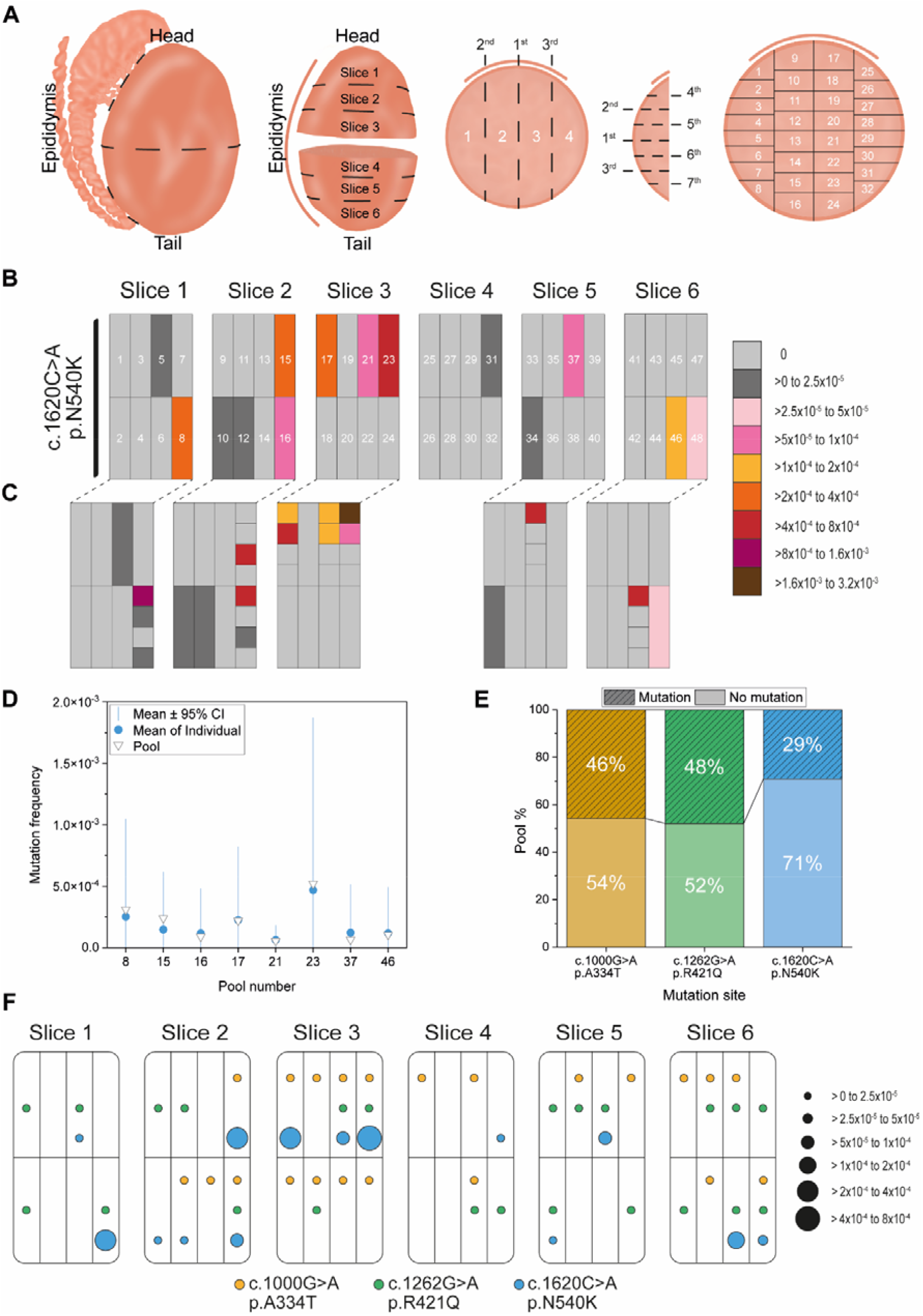
Testis data. **(A)** Testis cutting scheme strategy. The epididymis was removed, the testis was cut in half and fixated in 70% ethanol. Each half was further cut into 3 slices as shown. Each individual slice was cut into 4 vertical strips and 7 horizontal cuts. In sum, each slice was dissected into 32 individual pieces as shown above. **(B and C)** Mutation distribution of variant c.1620C>A in human testis of a 73-year-old donor. **(B)** Data analysis of individual pool samples (each consisting of 4 individual adjacent testis pieces). **(C)** In-depth analysis of “hot” pools (no. 8, 15, 16, 17, 21, 23, 37, and 46) with a mutation frequency above 5×10^−5^ to 1×10^−4^ (pink). **(D)** Mutation frequency accuracy test in eight different testis DNA pools for probe c.1620C>A (p.N540K) using two strategies: coarse vs fine pooling. Pool measurements (**▼**) represent the mutation frequency detected in the DNA of four testis pieces. The same testis pieces were measured individually and the respective mean (**○**) and 95% confidence intervals (CI) are represented. **(E)** Percentage of pools that harbor mutations for the three measured variants. **(F)** Mutation distribution of variants c.1000G>A, c.1262G>A, and c.1620C>A in the human testis of a 73-year-old donor. Data analysis of individual pool samples (each consisting of 4 individual adjacent testis pieces). Empty fields represent measurements with no mutations.

We adapted the testis micro-dissection technique commonly used in combination with single-molecule PCR screening (Choi et al., 2012, 2008; Qin et al., 2007) and employed ddPCR to quantify and study the spatial distribution of our target loci. Briefly, the human testis was dissected into 6 slices and further cut into 192 pieces (Figure 3A). We initially tested the resolution of a coarser pooling strategy. For this purpose, we compared two setups with the c.1620C>A variant: 1) Coarse pooling with 4 individual pieces (Figure 3B) vs 2) 32 individual pieces out of 192 pieces (Figure 3C).

Each pool measurement consisted of four adjacent testis pieces spatially organized according to the testis cutting scheme (Figure 3A). For the finer strategy, we remeasured the respective sub-pieces (4 pieces) of the coarse pools with VAF > 5×10^−5^ to better understand the contribution of the individual pieces to mutant counts in a fine vs coarse pooling strategy (48 pools vs. 192 pieces) (Figure 3B-D). We observed that the measured frequency using the coarse pooling matched the mean of the individual piece measurements and is within the respective standard deviation (Figure 3D). This data suggests that a coarse pooling strategy (48 pieces) renders sufficient resolution to distinguish sub-clonal expansion events in the testis.

We illustrated in Figure 3F the spatial distribution of our three selected loci with the dot size relative to the VAF (see Supplementary Table S6 for detailed frequencies). We observed that the c.1620C>A mutation is not uniformly distributed throughout the testis but instead is highly clustered with the piece harboring the largest VAF being adjacent to pieces with a much lower or absent frequency (Figure 3B-C, and F). In fact, there are a small number of pieces with frequencies that are orders of magnitude greater than the remaining pieces, and only 14/48 pools (29%, Figure 3E) harbored mutations. The average VAF for the whole testis was measured at 3.5×10^−5^ (Table 3); yet, the highest incidence pool had a frequency of 5.2×10^−4^ (Table 3) and the respective sub-piece at a frequency of 1.8×10^−3^ (Table S6).

**Table 3:** Screening of c.1000G>A, c.1262G>A, and c.1620C>A *FGFR3* mutations in testis DNA using the pooling strategy. IQR: Interquartile mutation frequency. Max Pool: Maximum mutation frequency in pooled samples. Max/IQR: Ratio of maximum mutation frequency to testis IQR mutation frequency. F<IQR: Fraction of pools with a mutation frequency lower than the testis IQR mutation frequency. P95: Fraction of pools necessary to include 95% of the mutants.

| Loci | n | Median VAF | IQR | Mean VAF | Standard Deviation | Max Pool | Max/IQR | F<IQR (%) | P95 (%) | Variant expansion behaviour |
| --- | --- | --- | --- | --- | --- | --- | --- | --- | --- | --- |
| c.1000G>A | 48 | 0 | $4.5 \times 10^{-6}$ | $3.4 \times 10^{-6}$ | $5.3 \times 10^{-6}$ | $2.3 \times 10^{-5}$ | 5.1 | 75.0 | 43.5 | Unknown |
| c.1262G>A | 48 | 0 | $6.0 \times 10^{-6}$ | $4.1 \times 10^{-6}$ | $5.7 \times 10^{-6}$ | $2.2 \times 10^{-5}$ | 3.7 | 72.9 | 45.5 | Unknown |
| c.1620C>A | 48 | 0 | $4.0 \times 10^{-6}$ | $3.5 \times 10^{-5}$ | $9.7 \times 10^{-5}$ | $5.2 \times 10^{-4}$ | 130 | 77.1 | 27.7 | Selective advantage |

Next, we introduced several summary statistics to quantify the amount of mutation clustering in the testis (Table 3). The ratio of the maximum pool VAF to the testis IQR (interquartile range) VAF (Max/IQR) is 130. In the case that mutations are uniformly distributed (no cluster formation), the Max/IQR should be closer to 1. We observed that most of the pools have a VAF below the IQR (e.g., for c.1620C>A the IQR is 4×10^−6^ and 77% of the pieces are below this value (F<IQR)). This data further supports that the variant c.1620C>A is not evenly distributed in the testis with some pieces having a very high VAF and others with no mutations. Similar mutational patterns have been previously reported in Apert syndrome (Choi et al., 2012, 2008; Maher et al., 2016; Qin et al., 2007), Noonan syndrome (Maher et al., 2018; Yoon et al., 2013), Multiple Endocrine Neoplasia Type 2B (MEN2B) (Choi et al., 2012), Thanatophoric Dysplasias (Goriely et al., 2009; Maher et al., 2016), Achondroplasia (Shinde et al., 2013), Pfeiffer syndrome (Maher et al., 2018, 2016), Hypochondroplasia, Crouzon, MEN2A, and Beare Stevenson syndromes (Maher et al., 2018) Finally, the fraction of the testis necessary to include 95% of the mutant counts (P95, 95% confidence) is ∼27.7%. Again, in a uniform mutation distribution in the testis, this fraction is expected to be close to 95%, i.e., only 5% of the pools do not harbor a mutation.

In contrast to the c.1620C>A variant, we observed that the c.1000G>A variant is found at a ten-fold lower average frequency of 3.4×10^−6^ in the whole testis and ∼46% of the pools (22/48) contained mutant cells (Figure 3E, Table 3, Supplementary Table S6). We found that the highest incidence pool (2.3×10^−5^) was only ∼5-fold higher than the interquartile mutation frequency (Max/IQR). Further, even though approximately 75% of the pools have a mutation frequency below 4.5×10^−6^ (F<IQR), 95% of our mutations were harbored in almost half of the pieces (∼44%). This minimal clustering suggests that mutant cells are tolerated only at low frequencies or they do not lead to such large clonal expansion events.

Similarly, in the third analyzed locus (c.1262G>A) the average VAF in the whole testis was as low as 4.1×10^−6^ (Table 3) with only half of the pools being positive for mutations (Figure 3E). The maximum VAF measured was 2.2×10^−5^ (Max Pool) which was only ∼3.7 fold higher in frequency compared to the testis IQR frequency (Max/IQR). This ratio is also closer to the expected proportion in the absence of clonal expansions. We further observed ∼73% of the pools having a lower mutation frequency than the IQR of 6×10^−6^ (F<IQR). Finally, our analysis shows a more uniform mutation distribution as ∼46% of the testis pools contained 95% of the mutants (expected P95 close to 95%, Table 3).

### *FGFR3* variants modify the receptor’s adaptor protein recruitment

Several studies have focused on the signaling effects of receptor tyrosine kinase (RTK) mutations (Foldynova-Trantirkova et al., 2012; He and Hristova, 2008; Li and Hristova, 2006; Meyers et al., 1995; Naski et al., 1996; Ornitz and Itoh, 2015; Toydemir et al., 2006; Wilkes et al., 1996). Growth factor availability is tightly controlled in the body via various mechanisms, as well as the interaction between fgfs and the extracellular matrix that influences the receptor affinity and diffusion of ligands through the tissue (Flaumenhaft et al., 1990; Ornitz, 2000). If an amino acid change is highly activating and independent of ligand binding, this might have the outcome of cells constantly receiving growth signals and lead to a growth advantage and clonal expansion typical of driver mutations.

Here, we sought to further investigate the activation of the studied *FGFR3* variants at the cellular level. For this, we used a protein micropatterning approach that allows the study of protein-protein interactions at the plasma membrane in living cells using total internal reflection fluorescence microscopy (Lanzerstorfer et al., 2014; Motsch et al., 2019; Schütz et al., 2017; Schwarzenbacher et al., 2008). In a recent study, we have demonstrated that this approach is quite robust for studying the activation level of FGFR3 when measuring the interaction with the downstream adaptor protein GRB2 (Hartl et al., 2022). Here, we co-expressed each of the different FGFR3 variants (labeled with mGFP) with GRB2 (labeled with mScarlet-I). In the plasma membrane, mGFP-FGFR3 is enriched and immobilized within certain areas that are defined by an antibody pattern on the cover slip. If FGFR3 is activated, the kinase domain gets phosphorylated and interacts with downstream signal adaptors such as GRB2-mScarlet-I and both fluorophore-tagged proteins co-localize within the same micropatterned areas. The degree of activation is proportional to the level of GRB2 recruited to active FGFR3, given here as the normalized mScarlet-I contrast, which relates the signal inside and outside the mGFP-FGFR3-enriched areas (for details see Methods section). Out of the eight analyzed variants (see Table S7 for summary statistics), two showed similar or lower activity than the WT receptor. The p.A334T (c.1000G>A) mutation located in the Ig-III extracellular domain of FGFR3 (Figure 4A) did not show a different receptor activity than the wild type (Figure 4B, Table 4 and Supplementary Table S8). In comparison, p.R621L (c.1862G>T) located in the kinase domain, specifically in the C-lobe of the split tyrosine kinase domain ((Farrell and Breeze, 2018); TK2 in Figure 4A) significantly inhibited the receptor activity.

**Table 4:** Summary of variant behavior in terms of the sperm VAF increasing with the donor’s age, percentage of donors with mutations in sperm, clustering in the testis, and activation of FGFR3 relative to the wild type and its response to ligand activation. AA: Amino Acid. CADD: Combined Annotation Dependent Depletion score. VAF: Variant Allele Frequency. TK: Tyrosine kinase. TM: Transmembrane. Ig-III: Immunoglobulin-like domain III. NA: Not Analyzed

| Significance (ClinVar) | Target site | AA | CADD | VAF | Sperm increase | Fraction of mutant sperm | Testis clustering | Fold FGFR3 activation | Activation state |
| --- | --- | --- | --- | --- | --- | --- | --- | --- | --- |
| Pathogenic | c.1620C>A | p.N540K | 24.4 | $1.1 \times 10^{-5}$ | Positive | 63% | Sub-clonal expansion | 1.6 | High and ligand independent |
| Pathogenic/<br>Uncertain | c.1537G>A | p.D513N | 23 | $9.4 \times 10^{-6}$ | No correlation | 60% | NA | 1.5 | High and ligand independent |
| Pathogenic | c.1172C>A | p.A391E | 22.5 | $8.6 \times 10^{-6}$ | No correlation | 32% | NA | 1.7 | High and ligand independent |
| Likely benign/<br>Uncertain | c.1879G>A | p.E627K | 31 | $1.5 \times 10^{-5}$ | No correlation | 89% | NA | 1.8 | High and ligand independent |
| Uncertain | c.1000G>A | p.A334T | 22.2 | $9.1 \times 10^{-5}$ | Positive<br>(non-significant) | 88% | Weak clustering | 1.1 | Physiologic |
| . | c.1262G>A | p.R421Q | 32 | $1.7 \times 10^{-5}$ | No correlation | 82% | Weak clustering | 2.1 | High and ligand independent |
| . | c.1862G>T | p.R621L | 29.3 | $2.7 \times 10^{-6}$ | No correlation | 34% | NA | 0.4 | Inhibited |
| . | c.952G>A | p.D318N | 22.3 | $2.2 \times 10^{-5}$ | Positive | 93% | NA | 1.5 | High and ligand independent |

**Figure 4:**
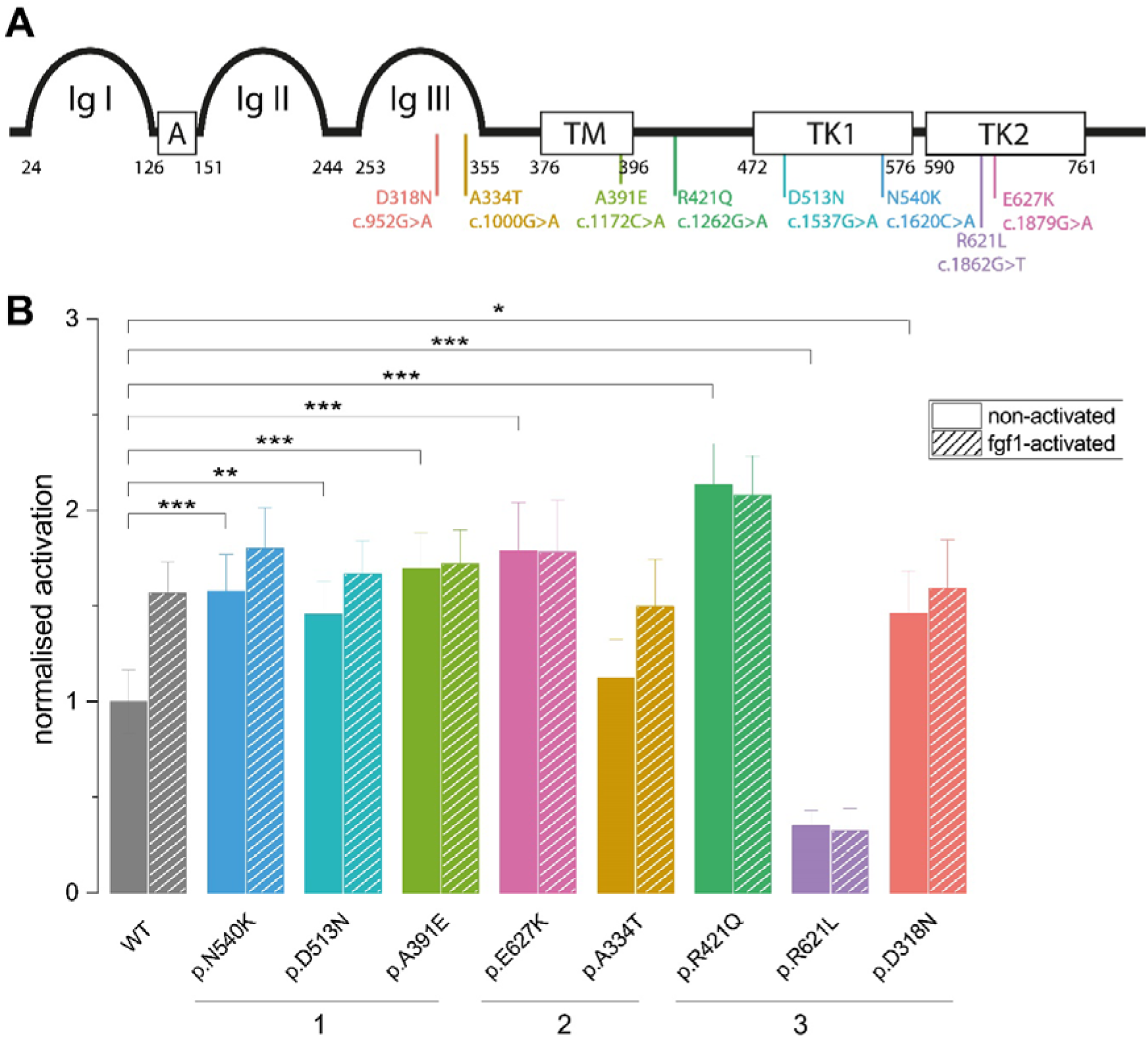
Signaling modulation by eight *de novo FGFR3* mutations in HeLa cells. **(A)** Schematic representation of the FGFR3 protein and the candidate variants. Mutations are indicated at their approximate location in the protein domains with their respective amino acid and nucleotide substitutions. Ig-like domains (Ig I-Ig III); acidic box (A), transmembrane domain (TM), and an intracellular split tyrosine kinase domain (TK1 and TK2). Numbers indicate the amino acid position of the respective domains. **(B)** The activation behavior of all candidate variants both in the non-activated and fgf1-stimulated state was normalized to the non-activated WT mScarlet-I contrast. The data is shown as normalized mean values with the respective standard error of the mean of a ratio as described in Supplementary Methods and Supplementary Table S7. The WT data set 1 (non-activated and fgf1-activated) was previously published in (Hartl et al., 2022). The number of cells measured for each variant is listed in Table S7. Only significant differences between the non-activated states for WT and the respective variants are indicated. The p-value (p) annotations are represented as follows: p≤0.05 (*), p≤0.01 (**), and p≤0.001 (***). Variant Significance Categories: 1) Pathogenic/Uncertain; 2) Likely benign/Uncertain; 3: Unclassified.

The remaining six variants lead to a significant increase in receptor signaling in the non-liganded state compared to the WT receptor (Figure 4B). These variants (c.1620C>A, p.N540K, TK1 domain; c.1537G>A, p.D513N, TK1 domain; c.1172C>A, p.A391E, TM domain; c.1879G>A, p.E627K, TK2 domain; c.1262G>A, p.R421Q and c.952G>A, p.D318N, Ig-III domain, locations shown in Figure 4A) presented a high activity with a 1.5 to about 2.5-fold increase in mScarlet-I contrast already in the absence of ligand which is comparable to the level of the ligand-activated WT receptor (Figure 4B). The difference in activation between WT and these six mutants was significant (see Table S8). Notably, only three out of these six variants have a reported deleterious score in ClinVar (p.N540K, p.D513N, p.A391E). However, variant p.R421Q located in the intracellular juxta membrane region of the receptor has the highest CADD score, yet has not been described in ClinVar. It also had the strongest promiscuous receptor activation (without ligand addition) compared to the other analyzed variants. None of the eight tested variants were further activated by the addition of ligand.

Overall, we observed that these *de novo* FGFR3 variants have different activation levels and therefore, recruit the adaptor protein GRB2 differently. Our data further suggests that these mutations modify the receptor signaling and therefore, affect the level of transmission throughout the male germline.

### Activating *FGFR3* substitutions do not equally expand in the male germline

The main characteristics of the variants (nucleotide and amino acid substitution, ClinVar significance, and CADD score) are summarized in Table 4 with the respective effect in the male germline reflected by several parameters: 1) the VAF, which represents the average mutation frequency in ∼177 donors; 2) an increase in VAF with the donor’s age representing testis-specific clonal expansion events; 3) evidence for clustering in the testis; and 4) the fraction of donors with measurable mutation frequencies reflecting recurrent, but independent *de novo* events.

With this sequence-function analysis, several important patterns can be observed. First, in general, a high and ligand-independent FGFR3 activation led to the accumulation of mutations in the male germline reflected by a VAF ∼ 10^−5^, and in some cases a positive correlation of VAF with age (Table 4). Notably, variant c.1000G>A (p.A334T) had the highest VAF, yet had an activation state similar to the wild type. Table 4 also shows that the levels of FGFR3 activation did not necessarily correlate with the VAF nor preclude strong sub-clonal expansion events in the testis or an increased number of mutations in older sperm donors. Second, FGFR3 activation resulted in a large fraction of donors with a measurable number of mutations (∼60-90%), except for c.1172C>A (p.A391E). This means that the mutations occurred independently and increased to measurable levels in a large sample size (young and old donors). Third, *in silico* annotations like the CADD score were not always accurate in predicting the biochemical effect of the mutation. Variants with similar CADD scores had different levels of ligand-independent activation. In one extreme case, variant c.1862G>T (p.R621L) was functionally very different (loss-of-function), albeit it had one of the highest CADD scores.

## Discussion

### Variants following a paternal-age effect

It has been hypothesized that gain-of-function substitutions that activate the RTK signaling pathway confer spermatogonial stem cells a selective advantage resulting in the sub-clonal expansion of mutant cells observed as testis-specific clusters and/or as a higher number of mutant sperm in older donors (Arnheim and Calabrese, 2009; Choi et al., 2008; Goriely et al., 2013; Goriely and Wilkie, 2012; Maher et al., 2014; Qin et al., 2007; Shinde et al., 2013; Yoon et al., 2013, 2009). In this work, we characterized the functional effect at the biochemical and cellular levels of eight different *FGFR3* variants. Five variants were reported in the literature to be associated with a congenital disorder and categorized to be of clinical significance by ClinVar (pathogenic or benign/uncertain). The other three have not been reported so far, yet were found at a higher frequency in sperm DNA (Salazar et al., 2022) and were predicted *in silico* to have a strong effect on modifying the protein structure (CADD score). In order to assess if the variant resulted in sub-clonal expansions in the male germline, we screened the VAF in two different cell types of the male germline: the spermatogonial stem cells and the differentiated sperm. Spermatogonial stem cells divide throughout the lifetime of the male germline and are thus a good target for expansions once hit by a driver mutation. We also used sperm to test whether mutations are passed on throughout the different maturation and differentiation stages of the male germline, including meiosis.

The sub-clonal expansion of mutant cells in the male germline predicts that mutations increase in frequency with age, also known as the paternal age effect (PAE). In this study, two out of the eight variants increased significantly in VAF with the age of the sperm donors: c.952G>A (p.D318N) and c.1620C>A (p.N540K). The first, a novel variant that has neither been characterized nor associated with disease, was measured at an average mutation frequency of 2.2×10^−5^, which is 2-fold higher in incidence than the second variant (Table 2, Figure 2). Further, this gain-of-function (activating) variant is viable (reported in gnomAD) and is found in sperm in a larger fraction of donors.

The c.1620C>A (p.N540K) variant showed the strongest increase in VAF with the age of the sperm donor (Figure 2). Recently, a study using duplex sequencing reported also a high VAF in sperm for this variant, together with a second substitution (c.1620C>G), resulting in the same amino acid change (p.N540K) (Salazar et al., 2022). Notably, the c.1620C>A (p.N540K) mutation has been described as the most prominent mutation causing Hypochondroplasia (Bellus et al., 1995; Ramaswami et al., 1998; Rousseau et al., 1996). Other congenital disorders such as Achondroplasia, Thanatophoric Dysplasia type I (Pannier et al., 2009), and Craniosynostosis associated with variant c.1620C>A (p.N540K) have also been reported in ClinVar. Moreover, c.1620C>A has been associated with several well-characterized PAE congenital disorders (e.g., ACH and TDI) and is the most common causative mutation for Hypochondroplasia (Bellus et al., 1995; Prinster et al., 1998; Ramaswami et al., 1998; Rousseau et al., 1996; Xue et al., 2014).

Further support to categorize the c.1620C>A as a canonical PAE-mutation that expands in the male germline comes from the testis dissection analysis. It has been observed in dissected testis or testis biopsies that certain mini-or micro-regions have accumulations of mutant spermatogonial stem cells (SSCs) several orders of magnitude higher than adjacent sites (Lim et al., 2012; Maher et al., 2016, 2018; Choi et al., 2008, 2012; Qin et al., 2007; Yoon et al., 2013; Goriely et al., 2009; Shinde et al., 2013) and are an indication of clonal expansion of a driver mutation in self-renewing SSCs over wild-type SrAp cells. We observed that the c.1620C>A was not uniformly distributed in the testis and presented the strongest clustering with large differences in VAF between adjacent pieces. A more recent study also documented clustering of the c.1620C>A mutation in one testis piece at mild frequencies (Maher et al., 2018). Maher *et al*., (2018) further reported a small cluster formation for the c.1620C>G mutation, which results in the same amino acid substitution p.N540K and is also associated with HCH. This together with our data suggests that both variants c.1620C>A and c.1620C>G with the same amino acid change might be an important driver mutation.

### What determines the sub-clonal expansion in the testis

Interestingly, two other analyzed variants c.1000G>A (p.A334T) and c.1262G>A (p.R421Q) showed only small, but unevenly distributed clusters, with mutants distributed in a larger proportion (46-48%) of the testis. In spite of this similar clustering behavior, these two variants showed very different activation states with the former showing no difference in receptor activation from the wild type; whereas, the latter had one of the highest constitutive activation states (∼2.5x activation). In particular, variant c.1000G>A (p.A334T) had the highest IQR frequency and a slight positive correlation of VAF with age (albeit not significant) suggesting a testis-specific sub-clonal expansion behavior. Variant p.R421Q (c.1262G>A) had an IQR VAF within the magnitude of most of our variants, but no increase in frequency with age.

Another interesting case is variant c.1172C>A (p.A391E). It is located in the transmembrane domain of the receptor and has been described to lead to constitutive receptor activation already in the absence of ligand (Chen et al., 2011; Mudumbi et al., 2013; Sarabipour and Hristova, 2016). We observed the same effect when measuring the GRB2 recruitment and we also showed that this variant cannot be further triggered by the addition of ligand. Although this mutation has one of the lowest pathogenicity scores of our candidate mutations, it is associated with Craniosynostosis and Crouzon Syndrome with Acanthosis Nigricans. Despite the association with germline disorders and the increased activating capabilities of the receptor, this variant has the second-lowest mutation frequency and the lowest number of positive donors (32%) of all the analyzed mutations. One could speculate that the activation of this variant still increases cell proliferation but might be tolerated only at very low levels. Spatial distribution analysis in the testis would be insightful in this regard.

### Phenotype of novel variants

Even though there are many *in silico* tools that predict the deleteriousness of a protein’s structure, the modification on the downstream signaling is unknown and can have different signaling outcomes. This was the case of the uncharacterized variants c.1262G>A (p.R421Q), c.1862G>T (p.R621L), and c.952G>A (p.D318N), all with high CADD scores, but very different signaling outcomes.

Variant c.1262G>A (p.R421Q) is located in the intracellular juxta membrane region of FGFR3 and has the highest predicted deleteriousness score, as well as the highest signaling activation detected with micropatterning, irrespective of fgf1 addition. Despite the fact, that c.1262G>A (p.R421Q) is not located in the catalytically active domain of the receptor, it resulted in the highest activation. Notably, this variant behaves similarly to the c.1879G>A (p.E627K) mutation in terms of activation and cellular effect (similar mutation frequency and number of positive donors in sperm). Yet, p.E627K variant is quite close to the catalytically active residue, in the C-lobe of the kinase domain, explaining the second highest activation. These findings indicate that the location of the mutation in the protein sequence is independent of the impact on the signaling output.

The second novel variant, c.1862G>T (p.R621L) showed a significantly decreased GRB2 recruitment compared to the WT receptor both in the non-liganded and the fgf1-activated state, similar to the established negative control (Hartl et al., 2022). This variant has never been described in the literature and has no associated syndromes. Although this mutation has one of the highest CADD scores, it has the lowest mutation frequency, and it is only found in 34% of donors. This variant actually decreases the receptor activity and therefore is possibly transmitted less in the germline. In line with our findings, a mutation in the same amino acid (p.R621H) has been reported to have a loss of function phenotype (Toydemir et al., 2006). The close proximity of the R621 amino acid to the invariant residue D618, which catalyzes phosphorylation within the conserved HRD motif (Farrell and Breeze, 2018), possibly explains the loss-of-function phenotype associated with mutations in this position.

The third novel variant c.952G>A (p.D318N) is located in the extracellular Ig-III domain of the receptor. The mutation leads to a low significant increase in GRB2 recruitment in the non-activated state as measured with micropatterning. The activation mechanisms might be explained by an altered affinity to fgf1 given that the ligand-binding in FGFR family members is regulated by the Ig-II and Ig-III domains as well as the linker region in between these domains (Ornitz and Itoh, 2015). Together with c.1620C>A (p.N540K), it is one of the only variants with a positive increase of VAF with age. It is detected in most of the tested donors with 93% of the sperm (n=86) being positive for this variant. This variant has not been associated with any syndromes and has a low predicted deleteriousness. Thus, we can speculate that the overactivation from this variant is well tolerated and might also clonally expand with age. Spatial distribution analysis in the testis of the p.D318N variant would further our knowledge of this hypothesis. Some parallels for c.952G>A (p.D318N) can be made with the mildly pathogenic c.1537G>A (p.D513N) mutation, associated with the Levy-Hollister syndrome. The Levy-Hollister syndrome has been reported as a unique case as it phenotypically differs from both gain-of-function associated congenital disorders (chondrodysplasias) and loss-of-function syndromes (skeletal over-growth and hearing loss) (Horton et al., 2007; Ornitz and Itoh, 2015; Toydemir et al., 2006). We measured for both variants one of the lowest activation levels in the non-liganded state and after the addition of fgf1. However, unlike p.D318N, the ddPCR results showed no significant enrichment in sperm with increasing age for variant p.D513N and was found in a lower number of donors (60% instead of 93%).

### Variants with unusually high *de novo* mutation rates

Precise ways of measuring *de novo* mutations rates developed in the past decade (trio sequencing (Campbell et al., 2012; Goldmann et al., 2016; Kong et al., 2012) and duplex sequencing (Abascal et al., 2021; Moore et al., 2021)) have shown that mutation rates depend on the local genetic context, the location within the genome, and the mutation type (reviewed in (Melamed et al., 2022)). However, these measurements represent mutations scattered randomly across the genome. Further, DNMs rate estimates in the germline are based on genome averages, instances of motifs (CpG sites), or complete genes (Campbell et al., 2012; Carlson et al., 2018; Goldmann et al., 2016; Kondrashov, 2003; Kong et al., 2012; Nachman and Crowell, 2000). The chance of encountering any particular mutation at a given specific site is very small and, as done in this study, only measurable with high accuracy methods that capture ultra-rare variants at single-mutation resolution by analyzing one specific nucleotide substitution within large populations of cells (>10^7^) in many individuals.

Our single-mutation resolution study showed that for six of the variants, 60-90% of the sperm donors harbored a measurable number of mutations (average VAF ranged between 1×10^−5^ to 9×10^−5^). This large proportion of individuals with DNMs of independent origin translates into a very high *de novo* mutation rate, much higher than expected from genome-wide averages. Several reasons could explain this trend: 1) the few measured mutant per donor counts are artifacts; 2) our variants are hypermutable sites in the *FGFR3* gene; 3) mutations occurred already throughout development in somatic tissue and are present at low levels in the male germline by a selective advantage. Given that for all the variants, we also measured donors without any mutations (sum of all screened genomes without a mutation (∼10^7^), the first explanation is not the most plausible one. A ‘hypermutable site’ explanation is also not very satisfying for missense coding variations.

A recent study that analyzed DNMs at single mutation resolution also identified a site with a very high DNM rate in the germline in the *HBB* gene resulting in the hemoglobin S (HbS). This high rate was mainly observed in the African population and it was hypothesized to be linked to a selective advantage by the mutation (protection against malaria by HbS) (Melamed et al., 2022). A similar hypothesis could be applied here. How can a selective advantage result in a high DNM rate? We suggest that mutations might also arise before the male germline becomes fully mature and have varying effects at different time points during development and throughout life. They can also appear post-zygotically, which would mean only a subset of cells would have such mutation (mosaic state) (Acuna-Hidalgo et al., 2016, 2015; Tiemann-Boege et al., 2021). During development, a series of random DNMs arise at any given time point in different cell lineages that change in number driven by different evolutionary processes. Some mutations changing the signaling might be linked to selection; whereby, one lineage is favored over another and produces more daughter cells, or sub-clonal expansions (Martincorena et al., 2018), as has been also observed in the testis for some of our *FGFR3* variants. The cluster size might also depend on the stochastic process of genetic drift (McFarland et al., 2014), the degree of selective advantage, and the interplay between cell growth and apoptosis (Arnheim and Calabrese, 2009; Choi et al., 2008; Goriely et al., 2013, 2005; Goriely and Wilkie, 2012; Maher et al., 2014; Qin et al., 2007; Yoon et al., 2013). The deleterious activating effect of the mutation might also have a negative outcome hindering cell fitness and decreasing the risk of transmission or might be tolerated only at very low levels (Tiemann-Boege et al., 2021). In contrast, milder activating mutations might have an overall selective growth advantage and accumulate more often at low levels, but in more cells (Goriely et al., 2013; Goriely and Wilkie, 2012). This selective advantage occurring already during development might also explain the behavior for variant c.1000G>A (p.A334T) having a physiological activation state, but observed at very high VAF in almost all donors.

## Conclusions

The present method allowed the in-depth analysis of millions of genomes of many individuals and can be used to investigate mutation rates of substitutions at single nucleotide resolution to understand the relationship between mutagenesis and selection of variants that change the signaling of the RTK pathway. We also showed the power of our approach in describing novel variants never reported before that have an effect at the cellular and biochemical level that also could have early- or late-onset consequences as a rare disease in offspring of older men.

## Methods

### Sample Collection

Sperm samples were collected from anonymous donors by the Kinderwunsch Klinik, MedCampus IV, Kepler Universitätklinikum, Linz according to the ethical approval (F1-11). Donors were aged from 23 to 59 years of age and mostly of European ancestry. Buccal samples were collected from an anonymous female donor aged 25 years old by swab collection. Cultured B-lymphocyte cells encoding the c.1620C>A variant in *FGFR3* (GM18666) were purchased from Coriell Institute (Camden, NJ).

### Sperm and Saliva DNA extraction

Genomic DNA from both sperm and saliva samples was extracted using the Puregene Core Kit A (QIAGEN, #1042601) according to the manufacturer’s instructions with minor modifications previously described in (Arbeithuber et al., 2015). In brief, DNA was extracted from a saliva swab or 25µl of sperm (∼ 10^6^ sperm cells) with the addition of 0.5µl of Proteinase K solution (20mg/µl) and 6µl of 1M DTT, followed by an overnight incubation step at 37°C. During DNA precipitation, 0.25µl of glycogen solution (QIAGEN, #1045724) were added. Vortexing steps were replaced by intensive manual shaking for 1 minute.

### Cultured cells DNA extraction

DNA extraction from cultured cells (GM18666) was carried out according to the manufacturer’s instructions using the Puregene Core Kit A (QIAGEN, #1042601). Initially, 200µl of cultured media (∼2×10^6^ harvested cells) were spun down for 5 seconds at 13,000g. The supernatant was discarded, and the cell pellet was vortexed and resuspended in the remaining 20µl of residual fluid. The cells were then resuspended at a high vortexing speed for 10 seconds in 300µl of Cell Lysis solution to promote cell lysis. Following this, 100µl of Protein precipitation solution was added and the sample was vortexed for an additional 20 seconds at high speed. The supernatant was transferred into a fresh tube containing 300µl of isopropanol and was gently inverted 50 times, followed by a centrifugation step at 13,000g for 1 minute. The supernatant was discarded, and the isolated DNA pellet was gently washed in 300µl of 70% ethanol and then centrifuged at 13,000g for 1 minute. The dried DNA pellet was then resuspended in 100µl of DNA hydration solution.

### Testis dissection

Testis (ID: NRD#ND10354) was acquired from the National Disease Research Interchange (NDRI, Philadelphia, PA) and procured by the same entity 5.7 hours after the time of death. Snap-frozen, post-mortem testis from a 73-year-old anonymous Caucasian donor with no precedents of alcohol, tobacco, or drug use was used in this study. The donor had no history of diabetes, chemotherapy, radiation, infectious diseases, or drug prescriptions.

In brief, the testis was initially thawed overnight on ice at 4°C. The epididymis was removed, and the testis was cut in half and fixated for 72 hours in 70% ethanol at 4°C. The testis was dissected into a total of 6 slices, each of them further cut into 32 pieces as previously described (Choi et al., 2008; Qin et al., 2007). For further information about the cutting strategy see Figure 3A.

### Testis DNA extraction

Testis DNA extraction was carried out for each individual testis piece using the NucleoMag Tissue Kit (Macherey-Nagel, #744300.1) according to the manufacturer’s conditions except for a few modifications. In a round bottom tube, up to 20mg of tissue immersed in 100µl of Buffer T1 and with the aid of a 5mm steel bead (QIAGEN, #69989) was homogenized in the TissueLyser (QIAGEN) at 25Hz for 1 minute and then briefly spun down. The steel bead was carefully removed and 100µl of Buffer T1 and 25µl Proteinase K solution (75mg/2.6mL) were added and mixed well prior to overnight tissue lysis at 37°C.

### Preparation of controls

As control measurements, we used buccal samples and wild-type (WT) plasmid (see “Genomic DNA plasmid” section in Supplementary Methods for details) as a guideline to set thresholds for the ddPCR data interpretation and as a proxy for the false-positive rate given by this technique. For site c.1537G>A, ∼250ng of *E. coli* carrier DNA was added to the 100,000 genome copies of WT plasmid to have a similar number of molecules and bulk DNA compared with the sperm DNA. Since no differences were observed (data not shown), *E. coli* DNA was not added in further experiments.

In order to verify the accuracy and dynamic range of our method, positive controls were prepared by using either commercially available cell lines containing the mutation of interest or by producing plasmids containing our mutation of interest by site-direct mutagenesis (see Supplementary Methods, Site-Directed Mutagenesis). These modified plasmids or cell lines were used in spike-in serial dilutions experiments (1:10 to 1:10,000; Figure 1B) with a total input of ∼20,000 genomes, containing both commercially available human genomic DNA (100ng/µl; ClonTech, #636401) and mutant DNA from either a cell line (c.1620C>A) or mutant plasmids for the remaining target sites herein characterized (primers used to produce the mutated plasmids are listed in Supplementary Table S9). Three biological replicates were run for each dilution step. Additionally, for the dilution step of 1:1000, 2 PCR reactions were set-up (total analyzed genomes: ∼40,000) and for the 1:10,000, 4 reactions were done (total of ∼80,000 genomes). Mutation sites c.952G>A and c.1000G>A were prepared on the pcDNA3.1 backbone containing the complete *FGFR3* ORF (Isoform IIIc). Mutations for sites c.1172C>A, c.1262G>A, c.1537G>A, c.1862G>T, and c.1879G>A were introduced in the pCR2.1 vector containing a genomic fragment of 1887 bp of *FGRF3* (chr4:1804338-chr4:1806225) – see Supplementary Methods-Genomic DNA Plasmid section and Supplementary Table S10.

### Droplet Digital PCR (ddPCR)

The ddPCR Assay design online tool (available at: https://www.bio-rad.com/digital-assays) was used to design target site-specific assays (see Supplementary Table S1-method). For the experimental set-up, standard ddPCR reaction conditions according to the manufacturer’s instructions (BioRad) were used. Individual ddPCR reactions were composed of 10µl of 10x SuperMix for Probes (no dUTP), 6.7µl of Nucleic acid free water, 2µl of genomic DNA (∼125ng/µl; ∼36,000 genomes/µl), 1µl of probes (900nM of each probe and 100nM of each primer), and 0.3µl of restriction enzyme (MseI or CviQI;10U/µl). Note that for three target sites (see Supplementary Table S11), 0.05µl of USER enzyme were added and incubated at 37°C for 15 minutes. For each sample, 4 reaction replicates were set up (total of ∼300,000 genomes).

Previous to the ddPCR droplet formation, we enhanced target template accessibility by fragmenting the genomic DNA (∼300 to <10kb) with the abovementioned restriction enzymes. In-ddPCR restriction digest was carried out by incubating the reaction mix at room temperature for 15 minutes and then transferred into the cartridge, along with 70µl of droplet generation oil for probes. The gasket was securely placed and the droplets were formed in the droplet generator. Approximately 43µl of the newly formed droplet solution were transferred into a ddPCR 96-well plate and then sealed at 180°C for 5 seconds in the PX1 PCR Plate Sealer (BioRad). PCR was carried out with the following conditions: 95°C for 10 minutes, 40 cycles of 94°C for 30 seconds, 53°C to 55°C (see Table S1-method for target-specific information) for 1 minute, and finally a one-time step of 98°C for 10 minutes. Note that a ramp rate of 2°C/sec was applied to each step and the lid was heated to 105°C. The PCR plate was transferred into the droplet reader for an end-point data analysis using QuantaSoft Analysis Pro Software (version 1.7.4; Bio-Rad Laboratories Inc.). We determined the detection thresholds using the positive and negative controls as a guideline. We used the Poisson corrected data points in this study.

### Micropatterning experiments

Micropatterning surface preparation, activation of patterned cells, total internal reflection fluorescence (TIRF) microscopy image acquisition, and microscopy data analysis were performed as previously described in (Hartl et al., 2022) and Supplementary Methods. Only for measurements of p.D318N, p.A334T, p.R621L, p.E627K and another set of WT no epoxy-coated coverslips were used. Glass slides (30mm diameter, 1.5mm height; Epredia, Portsmouth, NH, USA) were treated in a plasma cleaner (PDC-002 Plasma Cleaner Harrick Plasma, Ithaca, NY, USA) for 10 minutes followed by coating with AnteoBind™ Biosensor (AnteoTech, Brisbane, Queensland, Australia) for 60 minutes at RT to allow the streptavidin binding. Following this, a stamping procedure was performed as previously described.

### Statistical analysis

The Mann-Whitney-U and Kruskal-Wallis tests were used to test if there were differences in variant allele frequency (VAF) between the different sperm diagnosis groups. Spearman’s test was used to assess the correlation between VAF and the sperm count (Million sperms/mL) for each individual mutation (see Table S4). Spearman’s correlation test was applied to each sperm dataset to test the correlation between VAF and age. Comparison between the sperm age categories was subject to an initial Kruskal-Wallis test and for those with significant differences, the Mann-Whitney-U test was further used for categorical age comparison. Lastly, micropatterning data was subjected to one-way-ANOVA testing to compare the means of the obtained contrast values (see Table S8).

## Abbreviations

ACH: Achondroplasia
ddPCR: droplet digital Polymerase Chain Reaction
DNMs: *De novo* mutations
*FGFR3*: Fibroblast growth factor 3
HCH: Hypochondroplasia
ORF: Open Reading Frame
PAE: Paternal Age Effect
PZM: Post-zygotic Mosaic
SSCs: Spermatogonial stem cells
TDI: Thanatophoric Dysplasia Type I
TIRF: Total internal reflection fluorescence

## Disclosure Declaration

The authors declare no competing interests.

## Author’s contributions

S.M., I.H., V.B., A.Y., M.B., Y.S., and T.M. performed the experiments. S.M. and I.H. analyzed the data. I.T.-B. conceived the project and provided funding. S.M., I.H., and I.T.-B. wrote the manuscript. All authors read and approved the final manuscript.

## Acknowledgements

Funding for IH and VB was provided by the Doctoral College “NanoCell” from the Austrian Science Fund (FWFW1250), ES (V538-B26, FWF); and ITB (P30867000; FWF) and the European Regional Development Fund (REGGEN ATCZ207). We would like to thank Renato Salazar for the fruitful discussions and Philipp Hermann for the advice on statistical testing.

## Notes

### Competing Interest Statement

The authors have declared no competing interest.

